# Social media mining and photo-identification detects the shift of long-term seasonal foraging habitat for a juvenile loggerhead sea turtle

**DOI:** 10.1101/2022.02.28.482324

**Authors:** Kostas Papafitsoros

**Author notes:** Corresponding author: Kostas Papafitsoros.

## Abstract

Animal imagery uploaded on social media has been identified as an important tool in wildlife research and conservation and has been used in a variety of recent studies. Here a case study is presented, where a detailed analysis of social media content revealed a shift of a long-term foraging habitat for a juvenile loggerhead sea turtle (*Caretta caretta*) in Greece. In particular, this individual was a long term resident of Zakynthos island, Greece, from 2016 until 2020 (5 consecutive summer seasons), regularly foraging on a nearshore reef, with no observations of it being made during the 2021 season. However regular social media image mining combined with photo-identification, detected this individual foraging in the Gulf of Corinth in August 2021, more that 200km away from his previous foraging habitat. This case study (i) shows the possibility for juvenile loggerheads to shift their foraging site even after long term use, with implications in capture-mark-recapture studies and (ii) once more highlights the usefulness of social media mining and citizen science in diverse aspects of sea turtle studies.

## Introduction

The rise of social media and image sharing, combined with the widespread use of mobile phones and cameras has been identified as an important tool in conservation science (Dickinson *et al*., 2012; Di Minin *et al*., 2015; Toivonen *et al*., 2019). This is achieved either through citizen science with individuals directly contributing their images to social media platforms and dedicated web-based applications (Berger-Wolf *et al*., 2017; Giovos *et al*., 2019) or via systematic data mining from social media (automatic or manual search for focal animal imagery) (Giovos *et al*., 2018; Papafitsoros *et al*., 2021). In particular, with respect to sea turtles, citizen science has been exploited in many different ways during the last years, in a variety of studies. These range from determining foraging and nesting habitat distributions (Hudgins *et al*., 2017; Montagna *et al*., 2017; Baumbach *et al*., 2019; Casale *et al*., 2020; Hanna *et al*., 2021; Read and Jean, 2021), and evaluating tourism pressure (Papafitsoros *et al*., 2021) to documenting polyandry in loggerhead sea turtles (Papafitsoros *et al*., 2022). Typically in these type of studies photo-identification is also employed, exploiting the unique facial scale pattern of sea turtles, which is stable throughout their lifetime (Schofield *et al*., 2008; Carpentier *et al*., 2016), allowing for monitoring of specific individuals. Here we report yet another case in which the combination of social media, citizen science and photo-identification led to the detection of an event which is difficult to document with other means (e.g. flipper tagging, satellite tracking), that is the shift of a long term seasonal foraging habitat for a juvenile loggerhead sea turtle.

The main study area (i.e., initial foraging habitat) is situated in Laganas Bay, Zakynthos, Greece (37° 43’N, 20° 52’E). Laganas Bay hosts a major breeding site for the Mediterranean loggerhead sea turtles (*Caretta caretta*) (Margaritoulis *et al*., 2003), with on average 1218 nests being laid on 5.5km of nesting beach (1984-2009 average, Margaritoulis, 2005; Margaritoulis *et al*., 2011) and it is protected within the framework of the National Marine Park of Zakynthos, established in 1999. Satellite tracking and long term in-water surveys have revealed that Laganas Bay is also a foraging habitat for more that 40 resident sea turtles, mostly males and juveniles, (Papafitsoros and Schofield, 2016; Schofield *et al*., 2020; Papafitsoros *et al*., 2021). The island is also a popular summer tourist destination with over 850,000 visitors between May and October, and an established wildlife-watching industry, where tourists observe turtles via organised boat tours or independently (by swimming/snorkelling or hiring boats) (Schofield *et al*., 2015; Papafitsoros *et al*., 2021). As a result, there is a plethora of sea turtle imagery uploaded to social media by tourists every year. This imagery complements existing photo-identification based studies (Schofield *et al*., 2020), support studies that are related to wildlife watching itself but also can document elusive aspects of sea turtle ecology. For instance, a recent study that took place during the 2018-2019 seasons, and was based on the number of photographs of sea turtles of Zakynthos uploaded by tourists to social media (Instagram), revealed that the resident individuals are subject to high tourism pressure linked to high risk of propeller injuries and boat strikes (Papafitsoros *et al*., 2021). Another study also based on browsing online images of loggerhead sea turtles of Zakynthos led to the first direct documentation of polyandry in that species (Papafitsoros *et al*., 2022).

The purpose of the present study is to further investigate the capability of social media mining in sea turtle research. In particular, we are interested in examining whether it can be used for detecting sea turtle movements in large spatial scales, by searching the social media for images of loggerhead sea turtles of Zakynthos, that were however taken in a different location.

## Methods

We followed the methods described in Papafitsoros *et al*., 2021, which we briefly summarise here. We searched the Instagram platform for videos and photographs of sea turtles of Zakynthos, taken in other locations in the Mediterranean, via the use of key “hashtags” (“#”) related to the species (e.g. #loggerhead, #caretta). A complete list of the used hashtags can be found in the supplementary material of Papafitsoros *et al*., 2021. The search was performed on an at least weekly basis from 1 May to 31 October, 2021. If the facial scales of the turtle were clearly visible, we manually compared it to the ones belonged to an existing 22-year photoidentification database, containing over 1400 individuals of the sea turtles of Zakynthos (Schofield *et al*., 2020; Papafitsoros *et al*., 2021).

## Results

One Instagram photograph that showed a resident individual of Zakynthos (turtle ID: “t323”), in a location other than Zakynthos was detected, via the hashtag #carettacaretta, Figures 1 and 2. The photograph was uploaded in 10 August 2021 showing an underwater encounter that took place the previous day, 9 August 2021, in Koutros beach in the Gulf of Corinth (38° 20’ 59”N, 22° 37’ 50”E) approximately 200 kilometres away from Laganas Bay. The exact details of the encounter as well as further images were provided by the Instagram user who had uploaded the photograph. According to the existing photo-database this individual was a long term resident of Zakynthos with at least five consecutive years of presence in Laganas Bay (2016-2020), based on intermittent in-water observations restricted to the months June-October, as well as social media data mining (2018-2020) for the months April-November, Figure 3. Notably this individual was not observed in Zakynthos during 2021, neither via in-water surveys nor in other social media images. Its straight carapace length (SCL) was estimated around 55-60 cm based on side-to-side comparisons to other residents of the area whose length was known (Schofield *et al*., 2020), classifying it as an immature turtle based on minimum carapace sizes for mature turtles in the Mediterranean (65 cm SCL, Casale *et al*., 2005; Rees *et al*., 2013). The majority of its observations in Zakynthos during 2016-2020 had occurred in a nearshore reef, foraging on the sponge *Chondrilla nucula* (phylum Porifera, class Demospongiae) (Papafitsoros and Schofield, 2016; Papafitsoros *et al*., 2021). The turtle was also observed foraging next to the same species of sponges in Koutros beach, Figure 2, but verification of the exact prey could not be achieved due to the low resolution of the images.

**Figure 1.**
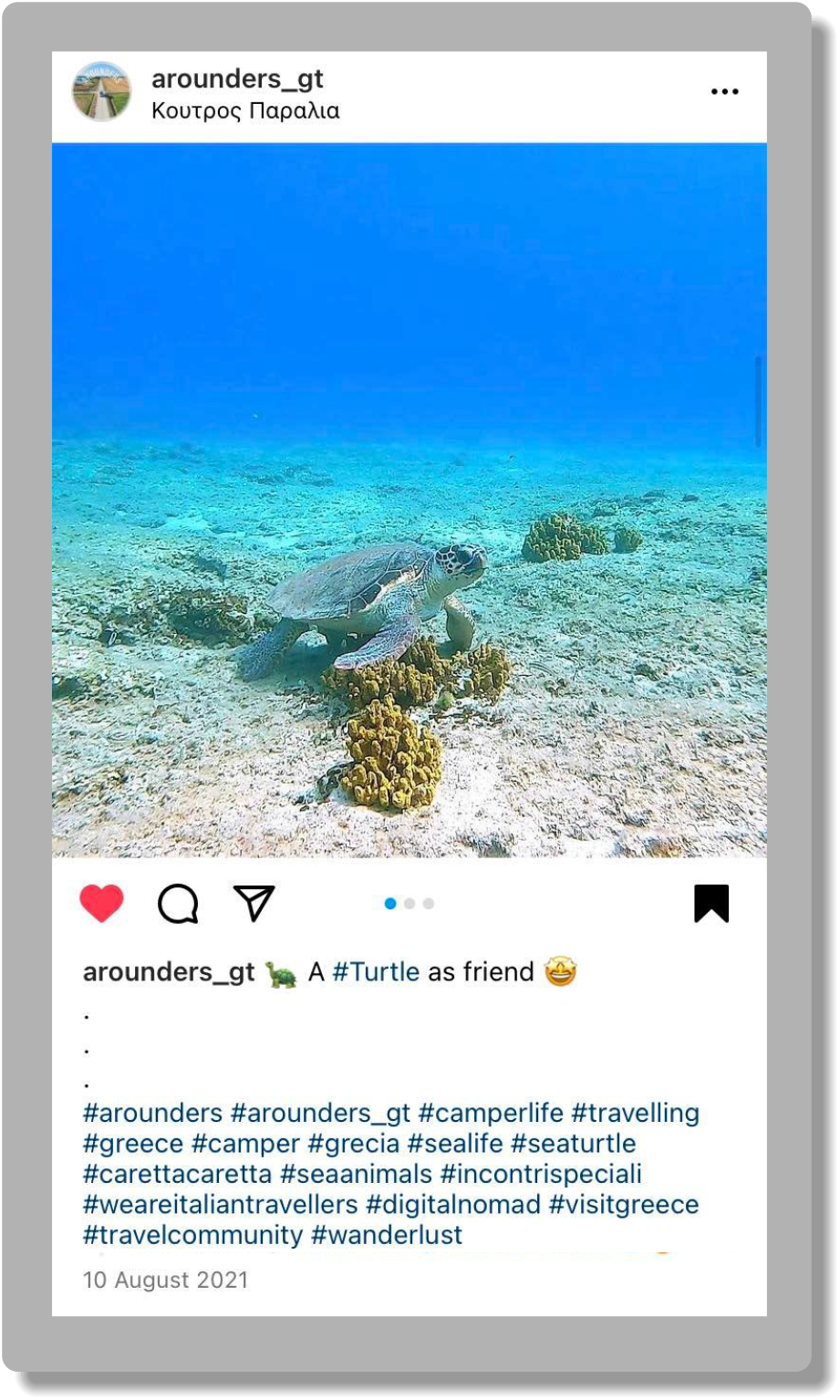
Original Instagram post uploaded in 10 August 2021, showing an underwater encounter with the loggerhead sea turtle “t323”, in Koutros beach, Gulf of Corinth, Photograph credits: Alessandra Villa & Luca Trespi (Instagram account: @arounders_gt)

**Figure 2.**
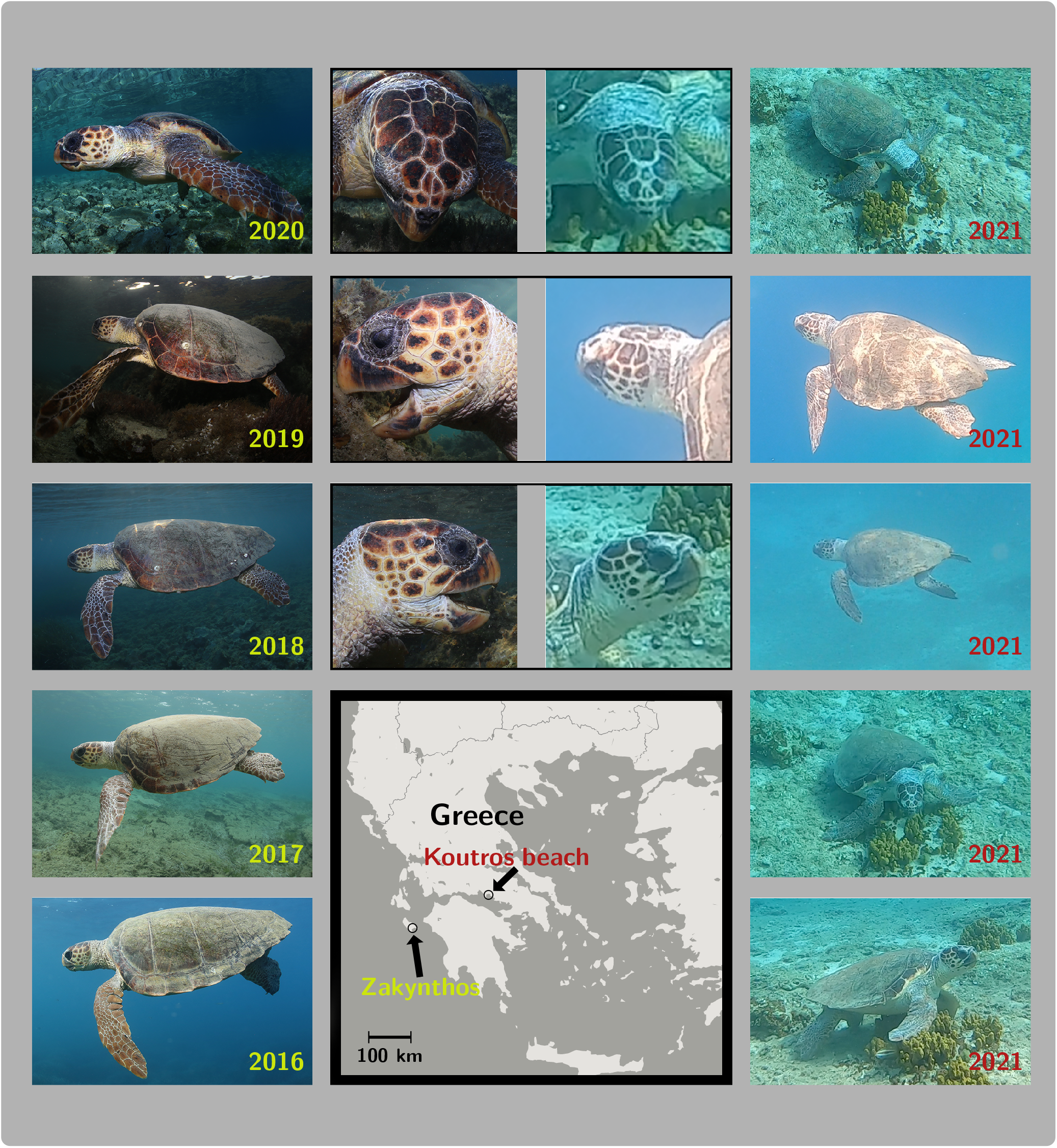
Left column: Photographs of the juvenile loggerhead turtle “t323” taken in Zakynthos, Greece, during the years 2016-2020. Right column: Photographs of the same individual uploaded on Instagram in 10 August 2021, showing an underwater encounter that took place the previous day, 9 August 2021, in Koutros beach, Gulf of Corinth, Greece. Middle top: Verification of the identity of the turtle via photo-identification based on top, left, and right facial scales. Middle bottom: Map of Greece, showing the location of Zakynthos and Koutros beach. Photograph credits: Kostas Papafitsoros (left), Alessandra Villa & Luca Trespi (right)

**Figure 3.**
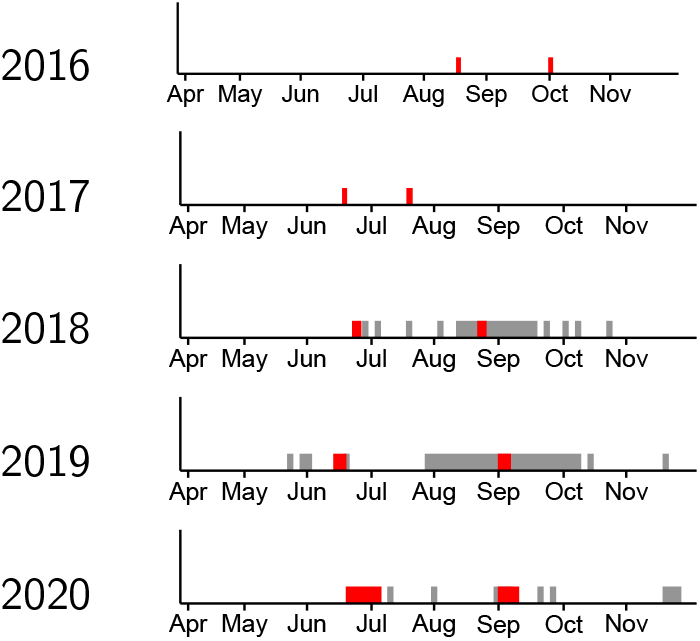
Timeline of observations of the individual “t323” in Zakynthos island for the years 2016-2020 showing (at least seasonal) residency for the months June-November. Red: direct in-water observations by the author. Grey: social media images (Instagram). No inspection of social media images took place in 2016-2017

## Discussion

Foraging site fidelity has been widely documented in sea turtles (Limpus *et al*., 1992; Broderick *et al*., 2007; Casale *et al*., 2012b; Rees *et al*., 2013; Shimada *et al*., 2020). Typically this refers to neritic foraging grounds as foraging site fidelity is not the norm for sea turtles in pelagic waters, where typically foraging is associated with continuous movements (Hawkes *et al*., 2006; Kobayashi *et al*., 2008; Rees *et al*., 2010). Foraging site fidelity is either confirmed via traditional capturemark-recapture (CMR) methods using flipper tags and/or satellite tracking. The latter is typically able to detect site fidelity of relative short time periods, typically only up to couple of years, constrained by the limited duration of satellite tracking devices (Hays *et al*., 2021). On the other hand traditional CMR studies on (neritic) foraging grounds are able to document individual histories spanning multiple decades (Rees *et al*., 2013; Shimada *et al*., 2020). Even in such long term studies, the detection of any foraging site shift can be quite challenging as it relies on either the existence of a similar CMR project in the new foraging site or on flipper tag reports by a third party, typically due to accidental bycatch in fisheries or death. For instance, from a selection of 175 flipper-tagged mature turtles recorded in foraging grounds in Australia (up to 29 year span), a foraging site shift (270 km) was recorded only for one female adult loggerhead (Shimada *et al*., 2020).

The rarity of this detection highlights the present case study even more, since not only a foraging site shift of more than 200km was detected for the individual “t323” but that occurred after at least 5 years of seasonal residency (June-November) in Zakynthos. However, given the zero survey effort and hence the absence of data supporting its presence in Zakynthos during the months December-May in 2016-2020, the case that this individual had already been using alternative foraging sites or even the newly detected site before 2021, cannot be excluded. Indeed, satellite tracking of two juvenile loggerheads originating from Drini Bay, Albania, (Snape *et al*., 2020), over one and two years respectively, revealed that these turtles departed to distant locations during autumn/winter months only to return to Drini Bay again in April/June. In contrast, the present study provides an additional case in which (at least) seasonal residency (summer months) can also be interrupted after many years. This has implications for CMR studies, since sea turtles that have not observed for many years are often “declared” dead in population survival analyses (Casale *et al*., 2007; Chaloupka and Limpus, 2002; Schofield *et al*., 2020). Here, a word of caution is provided, that this is not always the case and the possibility of a habitat shift even after a multi-year seasonal residence should be taken into account.

Sea turtles can change foraging habitats driven by a variety of factors such as prey availability (Dujon *et al*., 2018), water temperature (Casale *et al*., 2012a), predation risk (Heithaus *et al*., 2008), anthropogenic disturbances (Wright *et al*., 2020) as well interspecific interactions (Dujon *et al*., 2018). Given the high water temperatures during the summer months in both Zakynthos and the Gulf of Corinth, it is not likely that water temperature was a driving factor in our case. Moreover, the continuous presence of other turtles foraging on the same reef that this individual also occupied during 2016-2020 (author’s personal observations), suggests that prey remained available in that area during 2021 as well. Furthermore, predation risk in Zakynthos is extremely low for juvenile and adult sea turtles (Margaritoulis, 2005). In contrast, anthropogenic disturbances and interspecific interactions both occur in high rates in that reef (Papafitsoros and Schofield, 2016; Papafitsoros *et al*., 2021). In fact, the individual “t323” had being subjected to both immense and consistent tourist observation pressures (typically absent in the Gulf of Corinth) as well as aggressive interactions from the other foraging turtles in the area (Papafitsoros *et al*., 2021). Even though these could have been contributing factors for this habitat shift it is worth mentioning that some other turtles of Laganas Bay have remained (at least seasonal) residents for more than a decade, despite also being subjected to both high tourism pressures and aggressive interactions with other turtles (author’s personal observations).

The present study further highlights the possibility of conducting CMR studies through photoidentification. In fact, up to our current knowledge, this is the first time an identity of an alive sea turtle is recovered across such a large spatial scale (>200km), by the means of photo-identification only, without making use of flipper tags. Photo-identification relies on stable sea turtle external characteristics and thus can overcome the flipper tag-loss problem, a permanent point of concern for traditional CMR projects in both nesting and foraging grounds, (Pfaller *et al*., 2019). Photo-identification has already been employed in Zakynthos island for a variety of sea turtle studies, like survival analysis (Schofield *et al*., 2020), interspecific interactions in cleaning stations and foraging sites (Papafitsoros and Schofield, 2016; Schofield *et al*., 2017; Dujon *et al*., 2018), quantifying tourism pressure (Papafitsoros *et al*., 2021), as well as documenting polyandry (Papafitsoros *et al*., 2022). This study also took advantage of social media mining which together with dedicated citizen science platforms can help towards the collection of imaging data in large spatial and temporal scales and also retrospectively (Papafitsoros *et al*., 2021; Papafitsoros *et al*., 2022). Recent and continuous developments in automatic animal re-identification algorithms (Dunbar *et al*., 2021; Stewart *et al*., 2021) in combination with automatic mining for animal imagery in social media, open-data citizen science platforms (Berger-Wolf *et al*., 2017; Giovos *et al*., 2019), as well as advances in machine learning already facilitate these type of studies (Di Minin *et al*., 2018; Dujon and Schofield, 2019) and are expected to do so even more in the future.

## Acknowledgments

The author is grateful to Alessandra Villa and Luca Trespi (Instagram account: @arounders_gt) for providing all the relevant information regarding the encounter of this individual in Koutros beach, Gulf of Corinth, Greece.

## References

Baumbach, D.S., Anger, E.C., Collado, N.A., & Dunbar, S.G. (2019). Identifying sea turtle home ranges utilizing citizen-science data from novel web-based and smartphone GIS Applications. Chelonian Conserv. Biol. 18(2), 133–144. https://doi.org/10.2744/CCB-1355.1.

Berger-Wolf, T.Y., Rubenstein, D.I., Stewart, C.V., Holmberg, J.A., Parham, J., Menon, S., Crall, J., Van Oast, J., Kiciman, E., & Joppa, L. (2017). Wildbook: Crowdsourcing, computer vision, and data science for conservation. arXiv preprint arXiv:1710.08880.https://arxiv.org/abs/1710.08880.

Broderick, A.C., Coyne, M.S., Fuller, W.J., Glen, F., & Godley, B.J. (2007). Fidelity and overwintering of sea turtles. Proc. R. Soc. B 274(1617), 1533–1539. https://doi.org/10.1098/rspb.2007.0211.

Carpentier, A.S., Jean, C., Barret, M., Chassagneux, A., & Ciccione, S. (2016). Stability of facial scale patterns on green sea turtles Chelonia mydas over time: A validation for the use of a photo-identification method. J. Exp. Mar. Biol. Ecol. 476, 15–21. https://doi.org/10.1016/j.jembe.2015.12.003.

Casale, P., Affronte, M., Scarevelli, D., Lazar, B., Vallini, C., & Luschi, P. (2012a). Foraging grounds, movement patterns and habitat connectivity of juvenile loggerhead turtles (Caretta caretta) tracked from the Adriatic Sea. Mar. Biol. 159, 1527–1535. https://doi.org/10.1007/s00227-012-1937-2.

Casale, P., Broderick, A.C., Freggi, D., Mencacci, R., Fuller, W.J., Godley, B.J., & Luschi, P. (2012b). Long-term residence of juvenile loggerhead turtles to foraging grounds: a potential conservation hotspot in the Mediterranean. Aquat. Conserv. 22(2), 144–154. https://doi.org/10.1002/aqc.2222.

Casale, P., Ciccocioppo, A., Vagnoli, G., Rigoli, A., Freggi, D., Tolve, L., & Luschi, P. (2020). Citizen science helps assessing spatio-temporal distribution of sea turtles in foraging areas. Aquat. Conserv. 30(1), 123–130. https://doi.org/10.1002/aqc.3228.

Casale, P., Freggi, D., Basso, R., & Argano, R. (2005). Size at male maturity, sexing methods and adult sex ratio in loggerhead turtles (Caretta caretta) from Italian waters investigated through tail measurements. Herpetol. J. 15(3), 145–148.

Casale, P., Mazaris, A.D., Freggi, D., Basso, R., & Argano, R. (2007). Survival probabilities of loggerhead sea turtles (*Caretta caretta*) estimated from capture-mark-recapture data in the Mediterranean Sea. Sci. Mar. 71(2), 365–372. https://doi.org/10.3989/scimar.2007.71n2365.

Chaloupka, M. & Limpus, C. (2002). Survival probability estimates for the endangered loggerhead sea turtle resident in southern Great Barrier Reef waters. Mar. Biol. 140, 267–277. https://doi.org/10.1007/s002270100697.

Di Minin, E., Fink, C., Tenkanen, H., & Hiippala, T. (2018). Machine learning for tracking illegal wildlife trade on social media. Nat. Ecol. Evol. 2, 406–407. https://doi.org/10.1038/s41559-018-0466-x.

Di Minin, E., Tenkanen, H., & Toivonen, T. (2015). Prospects and challenges for social media data in conservation science. Front. Environ. Sci. 3, 63. http://dx.doi.org/10.3389/fenvs.2015.00063.

Dickinson, J.L., Shirk, J., Bonter, D., Bonney, R., Crain, R.L., Martin, J., Phillips, T., & Purcell, K. (2012). The current state of citizen science as a tool for ecological research and public engagement. Front. Ecol. Environ. 10(6), 291–297. https://doi.org/10.1890/110236.

Dujon, A.M. & Schofield, G. (2019). Importance of machine learning for enhancing ecological studies using information-rich imagery. Endanger. Species Res. 39, 91–104. https://doi.org/10.3354/esr00958.

Dujon, A.M., Schofield, G., Lester, R.E., Papafitsoros, K., & Hays, G.C. (2018). Complex movement patterns by foraging loggerhead sea turtles outside the breeding season identified using Argos-linked Fastloc-Global Positioning System. Mar. Ecol. 39(1), e12489. https://doi.org/10.1111/maec.12489.

Dunbar, S.G., Anger, E.C., Parham, J.R., Kingen, C., Wright, M.K., Hayes, C.T., Safi, S., Holmberg, J., Salinas, L., & Baumbach, D.S. (2021). HotSpotter: Using a computer-driven photo-id application to identify sea turtles. J. Exp. Mar. Biol. Ecol. 535, 151490. https://doi.org/10.1016/j.jembe.2020.151490.

Giovos, I., Keramidas, I., Antoniou, C., Deidun, A., Font, T., Kleitou, P., Lloret, J., Matić-Skoko, S., Said, A., Tiralongo, F., & Moutopoulos, D. K. (2018). Identifying recreational fisheries in the Mediterranean Sea through social media. Fish. Manag. Ecol. 25(4), 287–295. http://dx.doi.org/10.1111/fme.12293.

Giovos, I., Kleitou, P., Poursanidis, D., Batjakas, I., Bernardi, G., Crocetta, F., Doumpas, N., Kalogirou, S., Kampouris, T.E., Keramidas, I., Langeneck, J., Maximiadi, M., Mitsou, E., Stoilas, V.O, Tiralongo, F., Romanidis-Kyriakidis, G., Xentidis, N.J., Zenetos, A., & Katsanevakis, S. (2019). Citizen-science for monitoring marine invasions and stimulating public engagement: a case project from the eastern Mediterranean. Biol. Invasions 21(3), 3707–3721. https://doi.org/10.1007/s10530-019-02083-w.

Hanna, M.E., Chandler, E.M., Semmens, B.X., Eguchi, T., Lemons, G.E., & Seminoff, J.A. (2021). Citizen-Sourced Sightings and Underwater Photography Reveal Novel Insights About Green Sea Turtle Distribution and Ecology in Southern California. Front. Mar. Sci. 8. https://doi.org/10.3389/fmars.2021.671061.

Hawkes, L.A., Broderick, A.C., Coyne, M.S., Godfrey, M.H., Lopez-Jurado, L.F., Lopez-Suarez, P., Merino, S.E., Varo-Cruz, N., & Godley, B.J. (2006). Phenotypically Linked Dichotomy in Sea Turtle Foraging Requires Multiple Conservation Approaches. Curr. Biol. 16(10), 990–995. issn:0960-9822. https://doi.org/10.1016/j.cub.2006.03.063.

Hays, G.C., Laloë, J.-O., Rattray, A., & Esteban, N. (2021). Why do Argos satellite tags stop relaying data? Ecol. Evol. 11(11), 7093–7101. https://doi.org/10.1002/ece3.7558.

Heithaus, M.R., Wirsing, A.J, Thomson, J.A., & Burkholder, D.A. (2008). A review of lethal and non-lethal effects of predators on adult marine turtles. J. Exp. Mar. Biol. Ecol. 356(1), 43–51. https://doi.org/10.1016/j.jembe.2007.12.013.

Hudgins, J., Hudgins, E., Ali, K., & Mancini, A. (2017). Citizen science surveys elucidate key foraging and nesting habitat for two endangered marine turtle species within the Republic of Maldives. Herpetology Notes 10, 463–471.

Kobayashi, D.R., Polovina, J.J., Parker, D.M., Kamezaki, N., Cheng, I.J., Uchida, I., Dutton, P.H., & Balazs, G.H. (2008). Pelagic habitat characterization of loggerhead sea turtles, Caretta caretta, in the North Pacific Ocean (1997–2006): Insights from satellite tag tracking and remotely sensed data. J. Exp. Mar. Biol. Ecol. 356(1), 96–114. https://doi.org/10.1016/j.jembe.2007.12.019.

Limpus, C.J., Miller, J.D., Paramenter, C.J., Reimer, D., McLachlan, N., & Webb, R. (1992). Migration of green (Chelonia mydas) and loggerhead (Caretta caretta) turtles to and from eastern Australian rookeries. Wildl. Res. 19, 347–357. https://doi.org/10.1071/WR9920347.

Margaritoulis, D. (2005). Nesting activity and reproductive output of loggerhead sea turtles, Caretta caretta, over 19 Seasons (1984-2002) at Laganas Bay, Zakynthos, Greece: The largest rookery in the Mediterranean. Chelonian Conserv. Biol. 4(4), 916–929.

Margaritoulis, D., Argano, R., Baran, I., Bentivegna, F., Bradai, M. N., Caminas, J. A., Casale, P., De Metrio, G., Demetropoulos, A., Gerosa, G., Godley, B.J., Haddoud, D.A., Houghton, J., Laurent, L., & Lazar, B (2003). Loggerhead sea turtles. Ed. by A. B. Bolten & B. E. Witherington. Smithsonian Institution Press, 175–198.

Margaritoulis, D., Rees, A.F., & Riggal, T. (2011). Reproductive data of loggerhead turtles in Laganas Bay, Zakynthos island, Greece, 2003-2009. Marine Turtle Newsletter (131), 2–6.

Montagna, M., Taher, A.R., & Mancini, A. (2017). Combining citizen science and photo identification to monitor a key green turtle feeding ground in the southern Egyptian Red Sea. African Sea Turtle Newsletter 7, 8–15.

Papafitsoros, K., Dimitriadis, C., Mazaris, A.D., & Schofield, G. (2022). Photo-identification confirms polyandry in loggerhead sea turtles. Mar. Ecol. e12696. https://doi.org/10.1111/maec.12696.

Papafitsoros, K., Panagopoulou, A., & Schofield, G. (2021). Social media reveals consistently disproportionate tourism pressure on a threatened marine vertebrate. Anim. Conserv 24(4), 568–579. https://doi.org/10.1111/acv.12656.

Papafitsoros, K. & Schofield, G. (2016). Focal photograph surveys: Foraging resident male interactions and female interactions at fish-cleaning stations. Proceedings of the 36th Annual Symposium on Sea Turtle Biology and Conservation, Lima, Peru.

Pfaller, J.B., Williams, K.L., Frick, M.G., Shamblin, B.M., Nairn, C.J., & Girondot, M. (2019). Genetic determination of tag loss dynamics in nesting loggerhead turtles: a new chapter in “the tag loss problem”. Mar. Biol. 166(97). https://doi.org/10.1007/s00227-019-3545-x.

Read, T. & Jean, C. (2021). Using Social Media and Photo-Identification for Sea Turtles of New Caledonia. Marine Turtle Newsletter 162, 25–29.

Rees, A.F., Al Saady, S., Broderick, A.C., Coyne, M.S., Papathanasopoulou, N., & Godley, B.J. (2010). Behavioural polymorphism in one of the world’s largest populations of loggerhead sea turtles *Caretta caretta*. Mar. Ecol. Prog. Ser. 418, 201–212. https://doi.org/10.3354/meps08767.

Rees, A.F., Margaritoulis, D., Newman, R., Riggall, T.E., Tsaros, P., J.A., Zbinden, & Godley, B.J. (2013). Ecology of loggerhead marine turtles Caretta caretta in a neritic foraging habitat: movements, sex ratios and growth rates. Mar. Biol. 160(3), 519–529. https://doi.org/10.1007/s00227-012-2107-2.

Schofield, G., Katselidis, K.A., Dimopoulos, P., & Pantis, J.D. (2008). Investigating the viability of photo-identification as an objective tool to study endangered sea turtle populations. J. Exp. Mar. Biol. Ecol. 360(2), 103–108. https://doi.org/10.1016/j.jembe.2008.04.005.

Schofield, G., Klaassen, M., Papafitsoros, K., Lilley, M., Katselidis, K.A., & Hays, G.C. (2020). Long-term photo-id and satellite tracking reveal sex-biased survival linked to movements in an endangered species. Ecology 101 (7), e03027. https://doi.org/10.1002/ecy.3027.

Schofield, G., Papafitsoros, K., Haughey, R., & Katselidis, K. (2017). Aerial and underwater surveys reveal temporal variation in cleaning-station use by sea turtles at a temperate breeding area. Mar. Ecol. Prog. Ser. 575, 153–164. https://doi.org/10.3354/meps12193.

Schofield, G., Scott, R., Katselidis, K.A., Mazaris, A.D., & Hays, G.C. (2015). Quantifying wildlife-watching ecotourism intensity on an endangered marine vertebrate. Anim. Conserv. 18(6), 517–528. https://doi.org/10.1111/acv.12202.

Shimada, T., Limpus, C.J., Hamann, M., Bell, I., Esteban, N., Groom, R., & Hays, G.C. (2020). Fidelity to foraging sites after long migrations. J. Anim. Ecol. 89(4), 1008–1016. https://doi.org/10.1111/1365-2656.13157.

Snape, R.T.E., Schofield, G., & White, M. (2020). Delineating foraging grounds of a loggerhead turtle population through satellite tracking of juveniles. Aquat. Conserv. 30(7), 1476–1482. https://doi.org/10.1002/aqc.3302.

Stewart, C.V, Parham, J.R., Holmberg, J., & Berger-Wolf, T.Y. (2021). The Animal ID Problem: Continual Curation. arXiv preprint arXiv:2106.10377.https://arxiv.org/abs/2106.10377.

Toivonen, T., Heikinheimo, V., Fink, C., Hausmann, A., Hiippala, T., Järv, O., Tenkanen, H., & Di Minin, E. (2019). Social media data for conservation science: A methodological overview. Biol. Conserv. 233, 298–315. https://doi.org/10.1016/j.biocon.2019.01.023.

Wright, M. K., Baumbach, D.S., Collado, N., Safi, S. B., & Dunbar, S.G (2020). Influence of boat traffic on distribution and behavior of juvenile hawksbills foraging in a marine protected area in Roatán, Honduras. Ocean Coast. Manag. 198, 105379. https://doi.org/10.1016/j.ocecoaman.2020.105379.

